# PURE makes PURE: reconstitution of the PURE cell-free system from self-synthesized non-ribosomal proteins

**DOI:** 10.64898/2025.12.17.694911

**Authors:** Seyed Saeed Mottaghi, Sebastian J. Maerkl

## Abstract

Building a universal biochemical constructor, an autonomously self-replicating biochemical system, is a major challenge in synthetic biology. The PURE cell-free system is an ideal starting point for exploring self-regeneration, and its 36 non-ribosomal proteins constitute the primary macromolecular components that must be regenerated. Here, we demonstrate that the PURE system can be reconstituted from proteins synthesized by PURE itself. We first show that each of the 36 non-ribosomal proteins can be individually synthesized in PURE. We then purify the PURE synthesized proteins as pooled subsets and reconstitute a fully functional PURE system by combining the subsets. Finally, we show that all 36 non-ribosomal PURE proteins can be synthesized simultaneously in a single PURE reaction and, after purification, can reconstitute a functional PURE system. Together, these results establish that the non-ribosomal protein components of the PURE system can be self-regenerated, representing a critical step toward the realization of a universal biochemical constructor.

## Introduction

Bottom-up synthetic biology aims to construct biochemical systems that replicate the behavior of living cells by assembling non-living biological components [1–5]. One of the core properties of life is self-replication or autopoiesis [6], which requires the system to regenerate its molecular components from basic building blocks. Developing a universal biochemical constructor, a biochemical system capable of self-regeneration, is a major focus of current research in the field.

The Protein Synthesis Using Recombinant Elements (PURE) is a reconstituted cell-free transcription-translation system [7] that provides a suitable starting point for developing a universal biochemical constructor because it is defined and consists of a relatively small number of components. In order to exhibit self-replication, this system must regenerate its macromolecules, assuming that small molecules are supplied as building blocks and continuously fed into the reaction [8]. Most studies have so far focused on regeneration of tRNA [9–11], ribosomes [12–15], non-ribosomal proteins [8, 16–21], as well as DNA replication [22, 23]. Some efforts have been made towards integrating protein self-regeneration and DNA replication [24, 25]. In addition to macromolecules, amino acid synthesis in the PURE system has also been reported [26, 27].

Non-ribosomal proteins are key components in this context. Doerr et al. expressed 32 non-ribosomal PURE proteins in the PURE system from a single DNA template, and by analyzing the resulting expression products, they showed that a significant proportion of the synthesized proteins were truncated at the C-terminus due to impaired ribosomal processivity [16]. Other work focused on functional activity: a serial dilution approach for semi-continuous protein synthesis in bench-top experiments was developed by transferring a portion of the PURE reaction expressing proteins into a fresh ΔPURE reaction. This method enabled partial regeneration of some PURE proteins in the second reaction [19]. In another study, the essentiality of individual non-ribosomal PURE proteins and their functionality when expressed in the PURE system were assessed, and the results summarized in a table called the PUREiodic Table [18]. Lavickova et al. demonstrated sustained self-regeneration of seven aminoacyl-tRNA synthetases (AARSs) as well as 2 AARSs and T7 RNAP using a microfluidic chemostat device [8]. Most recently, Hagino et al. achieved regeneration of 20 AARSs in the PURE system for up to 20 cycles of serial dilution [17]. A recent preprint claims the functional synthesis of 30 non-ribosomal PURE proteins by localized synthesis from immobilized DNA templates [21].

The first step toward achieving non-ribosomal protein self-regeneration in the PURE system is to demonstrate that all proteins can be functionally expressed within that system. Although several studies have reported regeneration of groups of non-ribosomal PURE proteins or examined the functionality of sets of proteins expressed in PURE, no study has yet demonstrated whether the PURE system can be fully reconstituted from PURE expressed proteins. To this end, we aimed to synthesize all 36 non-ribosomal PURE proteins in PURE, purify them, reconstitute a new PURE system with these purified proteins, and evaluate the functionality of the reconstituted system.

## Results

To assess synthesis of non-ribosomal PURE proteins with PURE, we divided them into five subsets based on their function and concentration in the original PURE formulation: AARS1, AARS2, TLFs (translation factors excluding EF-Tu), EF-Tu, and Enzymes (Figure 1a). Each his-tagged PURE protein was expressed in a separate reaction using PUREfrex, which uses non-his-tagged versions of the PURE proteins, making it compatible with his-tag purification. Equal reaction volumes were used for all proteins in each subset except for EF-G, EF-Tu, and T7 RNAP, which were synthesized in larger volumes (Figure 1b). Subsequently, all proteins within a subset were pooled together and purified using magnetic beads. To visualize the expression of each protein, BODIPY-lysine-tRNA was used in separate reactions. After purification of each subset, buffer was exchanged using mini dialysis devices, and the samples were concentrated by ultra-filtration.

**Figure 1.**
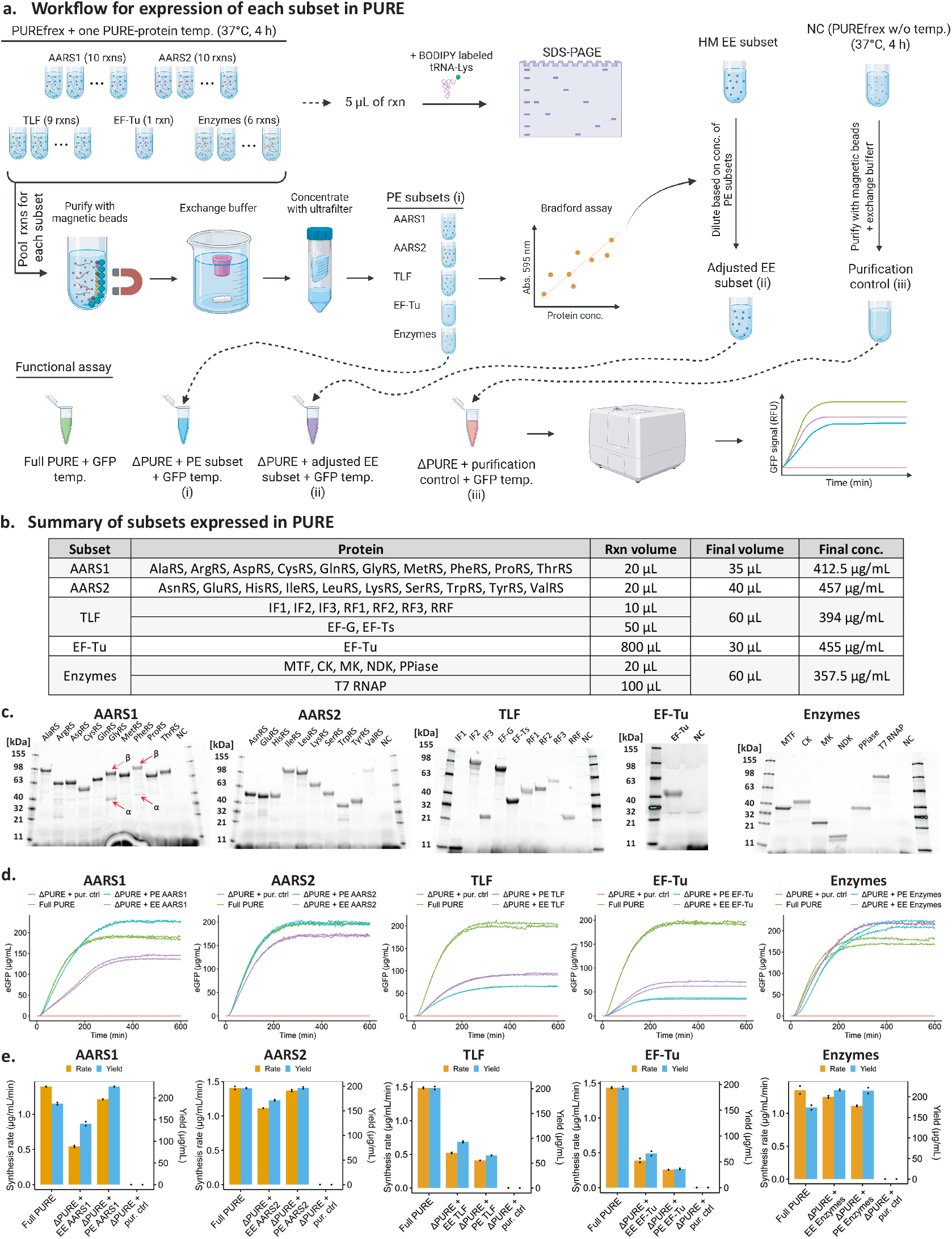
Synthesis of non-ribosomal PURE proteins by the PURE system, followed by functional evaluation. **a**, Workflow for the expression and purification of PURE proteins by the PURE system and their functional evaluation. Each his-tagged protein in a given subset was expressed using PUREfrex by adding the corresponding DNA template to the reaction and incubating at 37°C. Protein expression was confirmed by BODIPY-lysine fluorescence on SDS–PAGE. Reactions corresponding to each subset were pooled and purified using Ni-charged magnetic beads, followed by buffer exchange using mini dialysis devices and concentrated by ultra-filtration. For the functional assay, the purified subset was added to the corresponding ΔPURE reaction (a PURE reaction lacking the corresponding proteins) containing an eGFP template as a reporter for protein synthesis. Full PURE is included as the positive control, ΔPURE + purification control as the negative control, ΔPURE + EE subset as the concentration adjusted control (containing *E. coli* synthesized proteins at the same concentration as the PUREfrex-expressed subsets as determined by Bradford assay), and the ΔPURE + PE subset (PUREfrex expressed proteins). The fluorescence signal was measured using a plate reader. **b**, The table summarizes the PUREfrex reaction volumes used for each protein and the final volume and concentration obtained after purification of each pooled subset. **c**, SDS–PAGE analysis of individual PUREfrex reactions expressing PURE proteins. The gel shows the BODIPY-lysine fluorescence of each synthesized PURE protein. **d**, Functional assay result of each subset. The plot shows eGFP fluorescence in each reaction (n=2 for all conditions). **e**, Yield and rate of eGFP synthesis calculated from panel d.

The bands visualized via fluorescence using BODIPY-lysine incorporation on SDS-PAGE gels confirm the synthesis of each of the 36 non-ribosomal proteins (Figure 1c). All proteins were expressed at detectable levels. Some minor bands of lower molecular weight were observed, which could be attributed to truncated products. After purification, buffer exchange, and ultra-filtration of each subset we obtained total protein concentrations ranging from 356 to 457 µg/mL in volumes ranging from 30 to 60 µL (Figure 1b).

For the functional test, each purified subset was added to a ΔPURE reaction (a PURE reaction lacking the corresponding proteins) containing an eGFP template. These reactions were run alongside a negative control (ΔPURE supplemented with a purification control), a positive control (full PURE), and a concentration adjusted control. For the concentration adjusted control, we prepared the same subsets from individually expressed and purified proteins produced in *E. coli*. These proteins were added according to the original PURE formulation ratios, but the total concentration was adjusted to match those of the corresponding PURE expressed and purified subsets as determined by a Bradford assay.

Each purified protein subset, when added to a ΔPURE reaction background, resulted in eGFP expression, showing that each subset was able to restore functional protein synthesis (Figure 1d). None of the ΔPURE reactions supplemented with a purification control resulted in detectable eGFP synthesis indicating that no detectable carryover contamination from the original PUREfrex reaction occurred and that the omitted proteins as a group are essential for protein synthesis. Despite the AARS1 and AARS2 subsets being 4.7× and 3.5× lower concentration than the original formulation (Table S1) they exhibited yields and synthesis rates comparable to the full PURE positive control (Figure 1e), which is in concordance with previous results that indicate that AARSs are in excess concentration in the original PURE formulation [28]. The Enzymes subset concentration, which was 1.6-fold lower than in the original formulation, also resulted in similar rates and yields as the positive control and the concentration adjusted control. The TLF and EF-Tu subsets exhibited both lower yields and rates compared to the positive control and the concentration adjusted control. This is consistent with the critical concentration of these proteins in PURE (TLF and EF-Tu subsets were 5.4- and 4.6-fold less concentrated) [28]. The reduced yield and synthesis rates of the TLF and EF-TU subset compared to the positive control are likely primarily due to reduced and sub-optimal concentrations, as the difference to the concentration adjusted control is relatively minor. Nonetheless, both of these subsets performed less well than the concentration adjusted control which is most likely due to slightly reduced functionality of the PURE expressed protein, or possibly due to sub-optimal protein ratios in the case of the TLF subset. These results confirm that PURE is capable of functionally synthesizing its own non-ribosomal proteins. However, this conclusion cannot be confidently extended to every individual protein, as some PURE proteins are non-essential, and for others, protein impurities have been reported to interfere with individual dropout experiments [7, 17, 18, 21, 28].

After confirming the functionality of the individually PURE expressed, pooled, and purified protein subsets, the next step was to reconstitute a full PURE system by combining all five subsets together. To determine at what ratios these subsets should be combined, we titrated each subset in the corresponding ΔPURE reaction. AARS1, AARS2, and Enzymes subset concentrations were less critical; they reduced the reaction rate only if their concentration in the reaction dropped below 2.5%, 2.5%, and 5%, respectively. In contrast, EF-Tu and TLF concentrations were more limiting, and lowering them below 10% significantly reduced the reaction rate (Figures 2a, S1). Subsets were mixed according to the titration results and added to a ΔPURE reaction lacking all non-ribosomal proteins for a functional assay, along-side the same controls used previously (Figure 2b). As a concentration adjusted control, the corresponding subsets purified from *E. coli* with adjusted concentration were mixed in the same ratios as for the PUREfrex expressed subsets. The functional assay confirmed that the reconstituted PURE system is functional (Figure 2c) and exhibits protein synthesis yields and rates comparable to the concentration adjusted control (Figure 2d). The lower yield and rate compared to full PURE can therefore likely be mainly attributed to the lower concentration (7.7 times lower than full PURE, Table S1). This demonstrates that a functional PURE system can be fully reconstituted from PURE expressed proteins.

**Figure 2.**
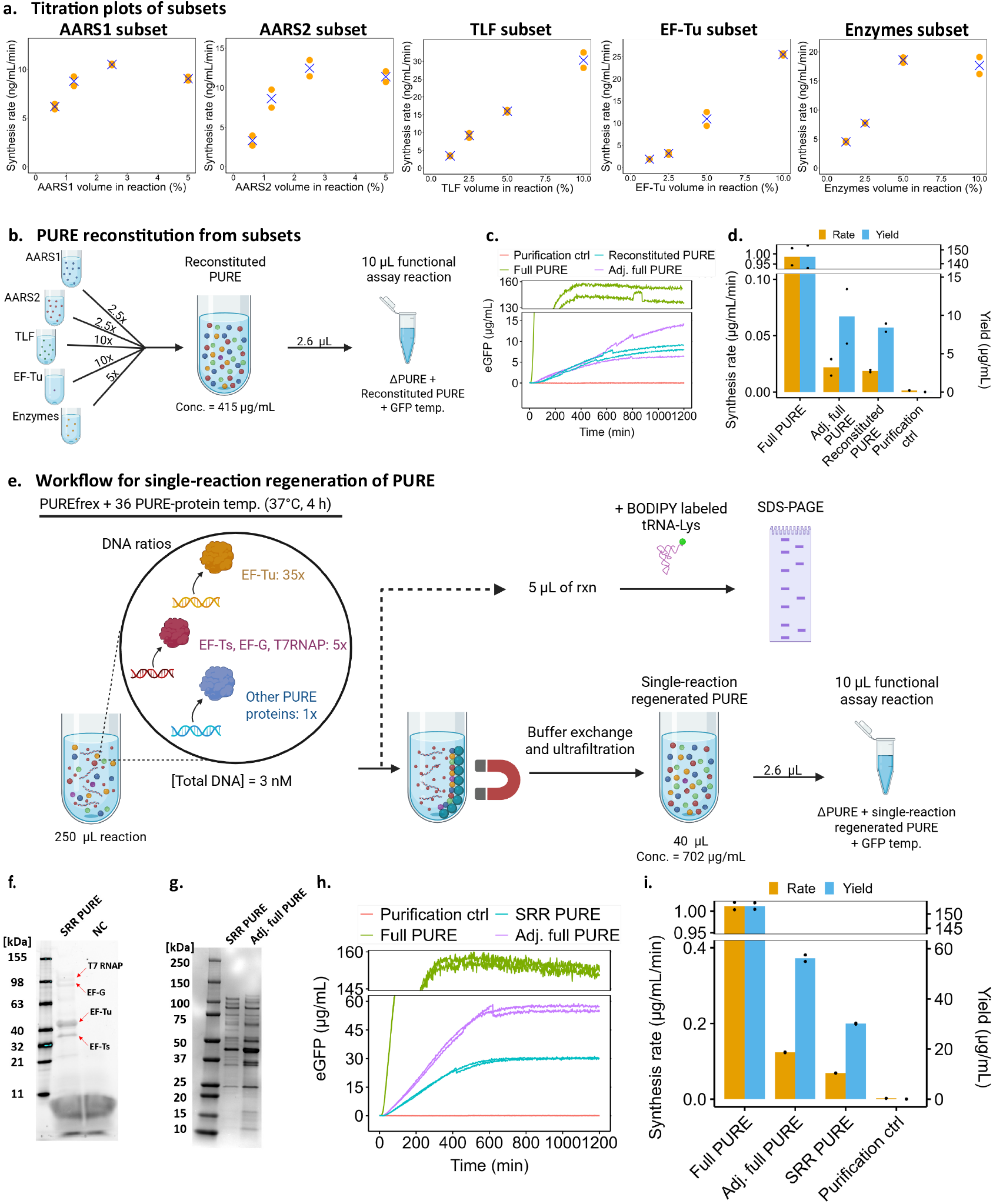
PURE fully reconstituted from PURE synthesized proteins. **a**, Titration of the different protein subsets in PURE. Each subset was titrated in its corresponding 10-fold diluted ΔPURE reaction (containing a 10-fold lower concentration of non-ribosomal proteins), and the optimal mixing ratio was determined based on the effect of each subset’s volume on the reaction rate. Orange dots show synthesis rate data points, and blue crosses represent their average. **b**, Reconstitution of PURE by combining functional subsets. After producing the five PURE synthesized protein subsets, a new PURE reaction was reconstituted by combining all subsets. The mixing ratios are indicated on the arrows. Subsequently, 2.6 µL of the reconstituted PURE protein solution was added to a ΔPURE reaction (lacking all non-ribosomal proteins) containing an eGFP template for a functional test. **c**, Functional assay result of reconstituted PURE assembled from the five subsets. The plot shows eGFP fluorescence generated in each reaction (*n* = 2 for all conditions). Full PURE is included as the positive control, ΔPURE supplemented with a purification control as the negative control, and the adjusted control was prepared from *E. coli* expressed proteins at equivalent concentrations, combined at the same ratios, and added to the ΔPURE reaction. **d**, Yield and rate of eGFP synthesis calculated from panel c. **e**, Single-reaction regeneration of PURE proteins in PURE. All 36 non-ribosomal proteins were expressed together in a single PURE reaction by including all DNA templates encoding the respective proteins. For EF-Tu, a 35-fold excess of DNA template was used, and for EF-Ts, EF-G, and T7 RNAP, a 5-fold excess was used. After incubation at 37°C, magnetic beads were used for purification, followed by buffer exchange and concentration. Subsequently, 2.6 µL of the single-reaction regenerated PURE was added to a ΔPURE reaction (lacking all non-ribosomal proteins) containing an eGFP template for a functional test. **f**, SDS–PAGE of single-reaction regenerated PURE proteins visualized by BODIPY-lysine fluorescence. **g**, Coomassie-stained SDS-PAGE of final obtained single-reaction regenerated PURE proteins next to concentration adjusted *E. coli* expressed full PURE. **h**, Functional assay result of single-reaction regenerated PURE. The plot shows eGFP fluorescence generated in each reaction (*n* = 2 for all conditions). Full PURE is included as the positive control, ΔPURE supplemented with a purification control as the negative control, and the adjusted control was prepared from homemade full PURE diluted to the same final total concentration and added to the ΔPURE reaction. **i**, Yield and rate of eGFP synthesis calculated from panel h.

Building on the successful reconstitution of a functional PURE system from PURE expressed proteins, we next sought to achieve simultaneous synthesis of all non-ribosomal PURE proteins in a single PUREfrex reaction, which more closely resembles the self - regeneration process required by a universal biochemical constructor. Synthesizing all proteins in a single reaction increases complexity, as expressing multiple genes in a single reaction leads to resource loading and competition [8], which can reduce the expression of proteins with lower expression levels relative to those expressed more efficiently. Furthermore, individual proteins are required at different concentrations. However, we simply assumed equal expression efficiencies for all gene templates and adjusted DNA ratios according to the desired protein concentrations in PURE. We therefore chose to use the following DNA concentrations: 35× for EF-Tu, 5× for T7 RNAP, EF-G, and EF-Ts, and 1× for the remaining PURE proteins. To slightly reduce resource loading, the total DNA concentration was lowered compared to the concentration used in the individual reactions above (3 nM instead of 5 nM). This was followed by the same purification, buffer exchange, concentration process, and functional test (Figure 2e). For the concentration adjusted control the homemade full PURE was diluted to the same concentration as the PURE expressed and purified total protein concentration.

The SDS-PAGE gel of BODIPY-lysine labeled products showed prominent expression bands corresponding to EF-Tu, T7 RNAP, EF-G, and EF-Ts, which had higher DNA template concentrations, whereas the remaining proteins were mostly below the detection limit (Figure 2f). A Coomassie-stained SDS-PAGE gel of the purified and concentrated single-reaction regenerated PURE system shows several additional bands, and an overall similar pattern to a concentration adjusted *E. coli*–expressed full PURE (Figure 2g).

The functional test confirms that single-reaction regenerated PURE is indeed functional (Figure 2h) and achieves yields comparable to those of the concentration adjusted control (Figure 2i). As before the lower yield and rate compared to full PURE can be mainly attributed to the lower total protein concentration (3.7-fold lower in the reaction, Table S1). In contrast, the reduced yield and rate relative to the concentration adjusted control may result from a less optimized formulation or reduced protein functionality. Interestingly, single-reaction regenerated PURE exhibits both higher yield and higher rate than reconstituted PURE, indicating that co-expression of multiple PURE proteins does not negatively affect their functionalities. The higher efficiency of single-reaction regenerated PURE may be due to its higher total protein concentration (702 vs. 415 µg/mL) or more favorable protein ratios. Thus, these results confirm that non-ribosomal PURE proteins can be synthesized in a single PURE reaction, purified, and used to reconstitute a new functional PURE system, representing an important step toward a universal biochemical constructor.

## Discussion

In this study we found that PURE is able to synthesize all 36 non-ribosomal proteins. We purified these proteins in five different pools and were able to recover PURE function using both individual pools as well as in a fully reconstituted PURE system in which all non-ribosomal proteins were synthesized by PURE. We took this approach one step further by demonstrating that we could express all 36 proteins in a single PURE reaction, and that this single pot reaction yielded sufficient protein to reconstitute a new PURE system.

One universal caveat of these types of studies demonstrating reconstitution of more than one protein at a time is the fact that the PURE system is not entirely pure. We and others have previously shown that several proteins in the PURE system are not essential when dropped out [17, 21, 28]. This appears to be mainly due to the fact that the ribosome fraction is often contaminated. Particularly, AARSs are often observed to be non-essential, although they theoretically should be. It is therefore a possibility that not all 36 non-ribosomal proteins need to be functionally expressed in order to give rise to a functional reconstituted system. This problem is exacerbated in experiments that require a seed concentration of the proteins being regenerated [21] as opposed to clean protein dropouts that ideally have been functionally validated as being essential [8].

Despite these caveats, the results obtained here support the notion that PURE can make PURE. Particularly since functional synthesis of EF-Tu and other ribosome factors, including IF3, EF-G, EF-Ts, was achieved, which constitute the majority of the PURE protein content and have previously been shown to be essential in our system [28]. This study represents an important step toward PURE self-regeneration, which is a crucial step toward building a universal biochemical constructor, and ultimately the construction of a synthetic cell.

## Methods

### Preparation of DNA templates

We used the same coding sequences as present in the original PURE plasmids (Addgene plasmids #124103–#124134, #118977, #124136, #118978, and #124138). Linear DNA templates with identical non-coding regions for all 36 PURE proteins as well as the eGFP linear template, were synthesized by Twist Bioscience, amplified by PCR using Q5 High-Fidelity 2X Master Mix (NEB, catalog no. M0492), and purified using a commercial kit (Zymo Research, catalog no. D4014). Figure S2 shows the final amplified and purified linear DNA templates for each PURE protein. The sequences of eGFP linear template and the 5’ and 3’ non-coding regions of PURE templates and the primers are listed in Table S2.

### Production of PURE proteins in *E. coli*

The same plasmids listed above were used for protein expression in *E. coli*. All PURE proteins were produced following the protocol described in Grasemann et al. [29]. After transforming the plasmids into either *E. coli* BL21(DE3) or M15 strains, single colonies were picked to inoculate precultures in a small volume of LB medium containing 100 µg/mL ampicillin (Amp), which were incubated overnight at 37°C and 260 rpm. The next day, cultures containing 100 µg/mL ampicillin were inoculated with 1% of the overnight cultures and incubated at 37°C and 260 rpm for 2 hours. Protein expression was then induced by the addition of isopropyl *β*-D-1-thiogalactopyranoside (IPTG) to a final concentration of 0.1 mM, followed by incubation for an additional 3 hours under the same conditions. Cells were harvested by centrifugation and stored at -80 °C. Buffers were prepared according to Table S3. For purification, the bacterial pellet was resuspended in buffer A and lysed by sonication. Cell debris was removed by centrifugation, and the resulting supernatant was loaded onto equilibrated IMAC Sepharose 6 Fast Flow beads (GE Healthcare, catalog no. GE17-0921-07). The column was washed with binding buffer and wash buffer containing 25 mM imidazole (23.75 mL of buffer A + 1.25 mL of buffer B), and the protein was eluted with elution buffer containing 450 mM imidazole (0.5 mL of buffer A + 4.5 mL of buffer B). The eluate was buffer-exchanged using a dialysis tubing cellulose membrane (Sigma-Aldrich, catalog no. D9652) against HT buffer overnight, and then against stock buffer for 3 hours. The final protein concentration was adjusted using a 0.5 mL Amicon Ultra Filter Unit (3 kDa MWCO; Millipore, catalog no. UFC5003).

After all PURE proteins were purified, homemade full PURE or different ΔPUREs were prepared by mixing the required proteins at their final concentrations, as in the original formulation.

### Expression of PURE system proteins in PURE

For the expression of PURE proteins in the PURE system, PUREfrex 2.0 (GeneFrontier, catalog no. GFK-PF201) was used following the manufacturer’s protocol. DNA templates for each protein were added to a final concentration of 5 nM. For single-reaction PURE regeneration, all DNA templates were added to the same reaction to a final total concentration of 3 nM. All templates were added at the same concentration, except for T7RNAP, EF-Ts, and EF-G, which were added at 5×, and EF-Tu, which was added at 35×. For the purification control, the PUREfrex master mix was prepared without any DNA template.

After preparing the reaction master mixes, 5 µL from each sample was taken, and 0.2 µL of BODIPY-lysine-tRNA (Promega, catalog no. L5001) was added. Reactions were incubated at 37°C for 4 hours to achieve maximum expression yield. The main samples were kept for purification, while the BODIPY-lysine labeled samples were run on SDS-PAGE. For preparation of BODIPY-lysine labeled samples for SDS-PAGE, 1 µL of 20× diluted RNase A (10 mg/mL; Thermo Scientific, catalog no. EN0531) was added to the reactions, followed by incubation at 37°C for 5 minutes. The subsequent steps were carried out according to the manufacturer’s protocol. A fluorescent protein standard (Invitrogen, catalog no. LC5928) was included on the SDS-PAGE as a reference.

### Purification of PURE expressed PURE subsets

After expression of the PURE proteins in individual reactions, all reactions corresponding to a given subset were pooled. The same buffers were used as those employed for the purification of PURE proteins from *E. coli*. For each 100 µL of total reaction, 50 µL of Ni-charged magnetic beads (GenScript, catalog no. L00295), corresponding to 12.5 µL of settled beads, were first washed twice with 500 µL of buffer A. The pooled samples were then added to the equilibrated beads and incubated at 4°C for 30 minutes on a rotisserie at 20 RPM. The single-reaction regenerated PURE was added to the beads in the same manner.

After incubation, the supernatant was removed using a magnetic rack (flow-through), and the beads were washed once with 750 µL of buffer A, followed by two washes with 750 µL of wash buffer (19 mL of buffer A + 1 mL of buffer B). Finally, 100 µL of elution buffer (0.5 mL of buffer A + 4.5 mL of buffer B) was added to the beads and incubated for 5 minutes on a rotisserie at 4°C. The supernatant containing the eluted target protein was collected. The elution step was then repeated once more, and the eluates were combined. For reactions with larger volumes, all quantities were scaled proportionally. Figure S3 shows the Coomassie-stained SDS-PAGE of samples from the different steps of the EF-Tu purification process. Since this protein was expressed and purified individually, its bands are clearly visualized by Coomassie staining.

### Buffer exchange and concentration of PURE expressed PURE subsets

For buffer exchange, samples were transferred into Slide-A-Lyzer™ MINI Dialysis Devices, 10K MWCO (Thermo Scientific, catalog no. 69574) and submerged in 100 mL of buffer HT. They were stirred gently at 150 RPM at 4°C overnight, followed by a second buffer exchange against 100 mL of stock buffer for 4 hours under the same conditions. After retrieving the samples, 0.5 mL Amicon Ultra Filter Units (Millipore; 3 kDa MWCO, catalog no. UFC5003, or 10 kDa MWCO, catalog no. UFC5010) were used to further concentrate the samples. The 3 kDa filters were used for the Enzymes and TLF subsets, while the 10 kDa filters were used for the AARS1, AARS2, and EF-Tu subsets. The purification control was not concentrated. The final obtained subsets were loaded on SDS-PAGE next to their adjusted controls, followed by Coomassie staining (Figure S4).

For protein concentration measurements, the Bradford assay was used. 2 µL of sample was mixed with 20 µL of Bradford dye (Bio-Rad, catalog no. 5000205). For the calibration curve, different concentrations of BSA (Sigma-Aldrich, catalog no. A3912) ranging from 50 µg/mL to 500 µg/mL were prepared in the same stock buffer and mixed with Bradford dye in the same way. The samples were incubated at room temperature for at least 5 minutes, and the absorbance was measured using a NanoDrop One spectrophotometer (Thermo Scientific). Protein concentrations were then calculated automatically by the instrument.

### eGFP synthesis reactions

All reactions had a total volume of 10 µL and were assembled using the energy solution (Solution I) and ribosomes (Solution III) from PUREfrex, as well as 10 nM of linear eGFP DNA template and the respective ΔPURE protein mixture. Ribosomes from PUREfrex were used because they have been reported to contain fewer protein impurities and produce lower background noise in dropout experiments [21]. For the functional test of subsets, 1 µL of the corresponding purified protein fraction was added to the ΔPURE reaction. For the functional test of full PUREs (either reconstituted PURE or single-reaction regenerated PURE), 2.6 µL was added. Because increasing the PURE protein volume increases the stock buffer added to the reaction, which has a negative effect on reaction yield. The added volume (2.6 µL) was determined by titrating different volumes of 20-fold diluted full PURE (containing a 20-fold lower concentration of non-ribosomal proteins) into PURE reactions. Above 2.6 µL, the reaction yield did not increase significantly; thus, this was determined to be the optimal quantity (Figure S5). In all functional tests, as the negative control, the same ΔPURE was supplemented with the same volume of the purification control to account for any non-ribosomal protein impurities from PUREfrex that may have non-specifically bound to the beads. As a positive control, homemade full PURE was used and supplemented with the same amount of stock buffer to compensate for the additional stock buffer present in the other samples. For adjusted controls, the ΔPURE reaction was supplemented with the same volume of *E. coli* expressed subset with adjusted concentration.

To determine the appropriate proportions for combining subsets to reconstitute PURE, each subset was titrated in the corresponding 10-fold diluted ΔPURE reaction (containing a 10-fold lower concentration of non-ribosomal proteins). The diluted reaction was used to better mimic the final reconstituted PURE system, in which protein concentrations are lower than those in the original PURE formulation. For these experiments, 10-fold diluted ΔPURE and serial dilutions of the subsets were prepared in the stock buffer. Subsequently, PURE reactions were assembled by adding the corresponding diluted ΔPURE and 1 µL of the diluted subset.

After preparing the samples, they were loaded into a 384-well black microplate with a transparent bottom (Corning, catalog no. 3544), and fluorescence was measured using a BioTek Synergy MX Multi-Mode Microplate Reader at 37°C (excitation: 488 nm; emission: 507 nm; gain: 70%).

### Calculation of yield and rate of eGFP synthesis

RFU values were converted to µg/mL based on a calibration curve. Different concentrations of recombinant eGFP (OriGene, catalog no. TP790050) were prepared by dilution in PBS and added to a PUREfrex reaction lacking a DNA template, as the eGFP signal is amplified in PURE compared to PBS. Then, fluorescence was measured under the same settings. Fluorescence values averaged over the second hour of measurement were used for curve fitting. The regression curve was constrained to pass through the mean RFU at 0 eGFP concentration. Experimental measurements were then converted using the resulting equation (Figure S6). Then to determine eGFP yield, converted values were averaged over the final hour of measurement. To calculate the synthesis rate, the slope was determined for each 60-minute interval using linear regression, and the maximum value was defined as the reaction rate.

## Author Contribution

S.S.M. performed experiments. S.S.M. and S.J.M. designed experiments, analyzed data, and wrote the manuscript.

## Acknowledgements

The authors would like to express their gratitude to Dr. Fanjun Li, Amogh Kumar Baranwal, and Pao-Wan Lee for helpful input and comments. We also thank Dr. Ragunathan Bava Ganesh for providing some of the purified PURE protein components and linear DNA templates. This work was supported by a Swiss National Science Foundation MINT grant (200020 214843). Graphical schematic representations were created with BioRender.com.

## References

[1] Guindani, C., da Silva, L. C., Cao, S., Ivanov, T. & Landfester, K. Synthetic cells: from simple bio-inspired modules to sophisticated integrated systems. Angewandte Chemie 134, e202110855 (2022).

[2] Hirschi, S., Ward, T. R., Meier, W. P., Muller, D. J. & Fotiadis, D. Synthetic biology: bottom-up assembly of molecular systems. Chemical Reviews 122, 16294–16328 (2022).

[3] Ivanov, I. et al. Bottom-up synthesis of artificial cells: recent highlights and future challenges. Annual Review of Chemical and Biomolecular Engineering 12, 287–308 (2021).

[4] Schwille, P. et al. Maxsynbio: avenues towards creating cells from the bottom up. Angewandte Chemie International Edition 57, 13382–13392 (2018).

[5] Laohakunakorn, N. et al. Bottom-up construction of complex biomolecular systems with cell-free synthetic biology. Frontiers in bioengineering and biotechnology 8, 213 (2020).

[6] Maerkl, S. J. On biochemical constructors and synthetic cells. Interface Focus 13, 20230014 (2023).

[7] Shimizu, Y. et al. Cell-free translation reconstituted with purified components. Nature biotechnology 19, 751–755 (2001).

[8] Lavickova, B., Laohakunakorn, N. & Maerkl, S. J. A partially self-regenerating synthetic cell. Nature Communications 11, 6340 (2020).

[9] Hibi, K. et al. Reconstituted cell-free protein synthesis using in vitro transcribed trnas. Communications Biology 3, 350 (2020).

[10] Li, F., Baranwal, A. K. & Maerkl, S. J. Continuous in situ synthesis of a complete set of trnas sustains steady-state translation in a recombinant cell-free system. Nature Communications 16, 6212 (2025).

[11] Miyachi, R., Masuda, K., Shimizu, Y. & Ichihashi, N. Simultaneous in vitro expression of minimal 21 transfer rnas by trna array method. Nature Communications 16, 7418 (2025).

[12] Jewett, M. C., Fritz, B. R., Timmerman, L. E. & Church, G. M. In vitro integration of ribosomal rna synthesis, ribosome assembly, and translation. Molecular systems biology 9, 678 (2013).

[13] Li, J. et al. Cogenerating synthetic parts toward a self-replicating system. ACS synthetic biology 6, 1327–1336 (2017).

[14] Shimojo, M. et al. In vitro reconstitution of functional small ribosomal subunit assembly for comprehensive analysis of ribosomal elements in e. coli. Communications Biology 3, 142 (2020).

[15] Aoyama, R. et al. In vitro reconstitution of the escherichia coli 70s ribosome with a full set of recombinant ribosomal proteins. J. Biochem. 171, 227–237 (2022).

[16] Doerr, A., Foschepoth, D., Forster, A. C. & Danelon, C. In vitro synthesis of 32 translation-factor proteins from a single template reveals impaired ribosomal processivity. Scientific Reports 11, 1898 (2021).

[17] Hagino, K., Masuda, K., Shimizu, Y. & Ichihashi, N. Sustainable regeneration of 20 aminoacyl-trna synthetases in a reconstituted system toward self-synthesizing artificial systems. Science Advances 11, eadt6269 (2025).

[18] Wei, E. & Endy, D. Experimental tests of functional molecular regeneration via a standard framework for coordinating synthetic cell building. BioRxiv 2021–03 (2021).

[19] Libicher, K. & Mutschler, H. Probing self-regeneration of essential protein factors required for in vitro translation activity by serial transfer. Chemical Communications 56, 15426–15429 (2020).

[20] Awai, T., Ichihashi, N. & Yomo, T. Activities of 20 aminoacyl-trna synthetases expressed in a reconstituted translation system in escherichia coli. Biochemistry and biophysics reports 3, 140–143 (2015).

[21] Schwarz-Schilling, M. et al. Autonomous biogenesis of the entire protein translation machinery excluding ribosomes. bioRxiv 2024–10 (2024).

[22] Sakatani, Y., Yomo, T. & Ichihashi, N. Self-replication of circular dna by a self-encoded dna polymerase through rolling-circle replication and recombination. Scientific Reports 8, 13089 (2018).

[23] Van Nies, P. et al. Self-replication of dna by its encoded proteins in liposome-based synthetic cells. Nature communications 9, 1583 (2018).

[24] Libicher, K., Hornberger, R., Heymann, M. & Mutschler, H. In vitro self-replication and multicistronic expression of large synthetic genomes. Nature Communications 11, 904 (2020).

[25] Hagino, K. & Ichihashi, N. In vitro transcription/translation-coupled dna replication through partial regeneration of 20 aminoacyl-trna synthetases. ACS Synthetic Biology 12, 1252–1263 (2023).

[26] Nishikawa, S. et al. Amino acid self-regenerating cell-free protein synthesis system that feeds on pla plastics, co2, ammonium, and α-ketoglutarate. ACS Catalysis 14, 7696–7706 (2024).

[27] Giaveri, S. et al. Integrated translation and metabolism in a partially self-synthesizing biochemical network. Science 385, 174–178 (2024).

[28] Ganesh, R. B. & Maerkl, S. J. Towards self-regeneration: Exploring the limits of protein synthesis in the protein synthesis using recombinant elements (pure) cell-free transcription–translation system. ACS Synthetic Biology 13, 2555–2566 (2024).

[29] Grasemann, L., Lavickova, B., Elizondo-Cantú, M. C. & Maerkl, S. J. Onepot pure cell-free system. Journal of Visualized Experiments 23 (2021).

