## Supplementary Information for "PURE makes PURE: reconstitution of the PURE cell-free system from self-synthesized non-ribosomal proteins"

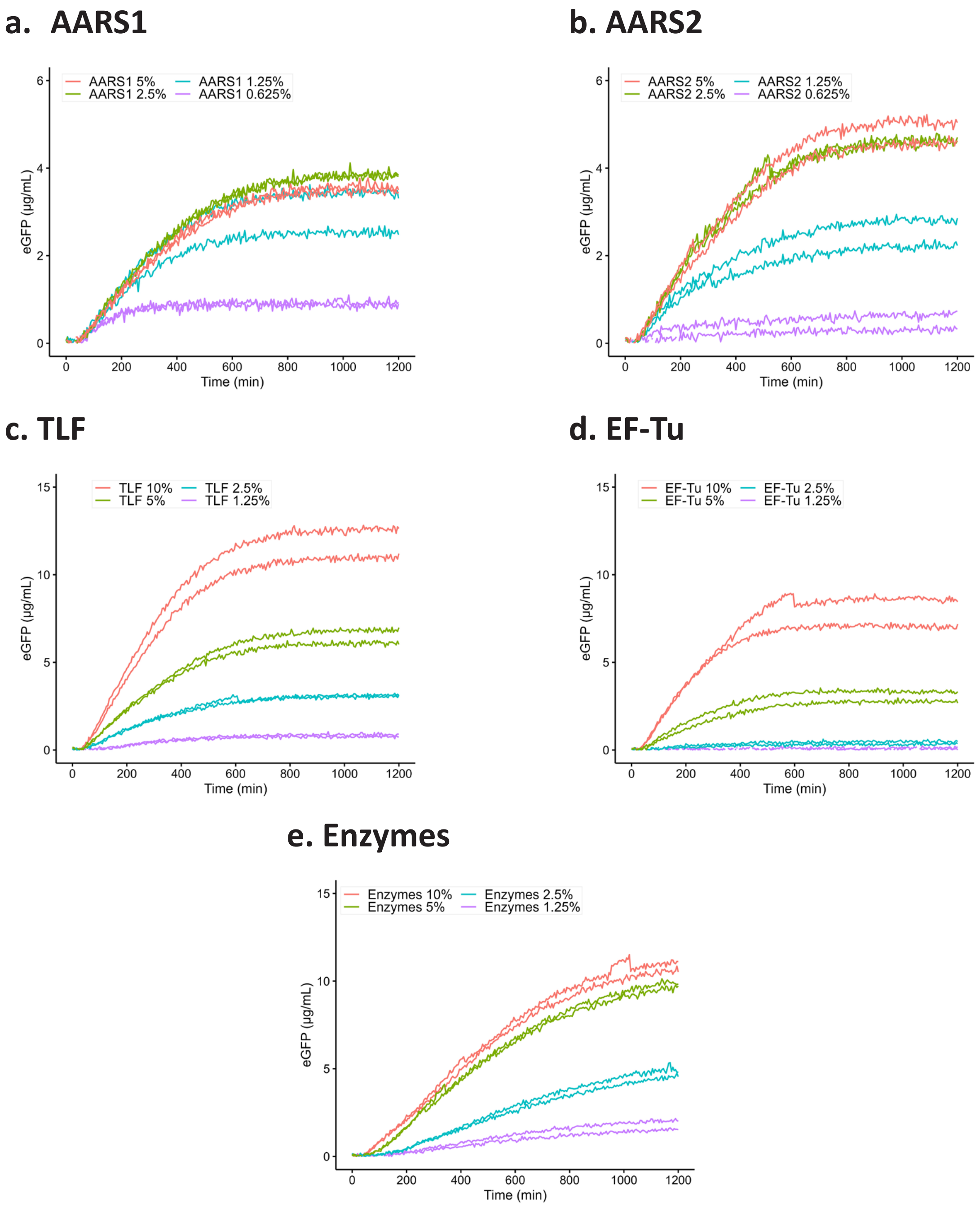


Figure S1: **Titration plots of PURE expressed protein subsets.** (a–e) Serial dilutions of each subset were prepared and added to the corresponding 10-fold diluted $\Delta$PURE reaction containing an eGFP template, and fluorescence was measured over time. The rates in the titration plots in Figure 2a are based on these results.

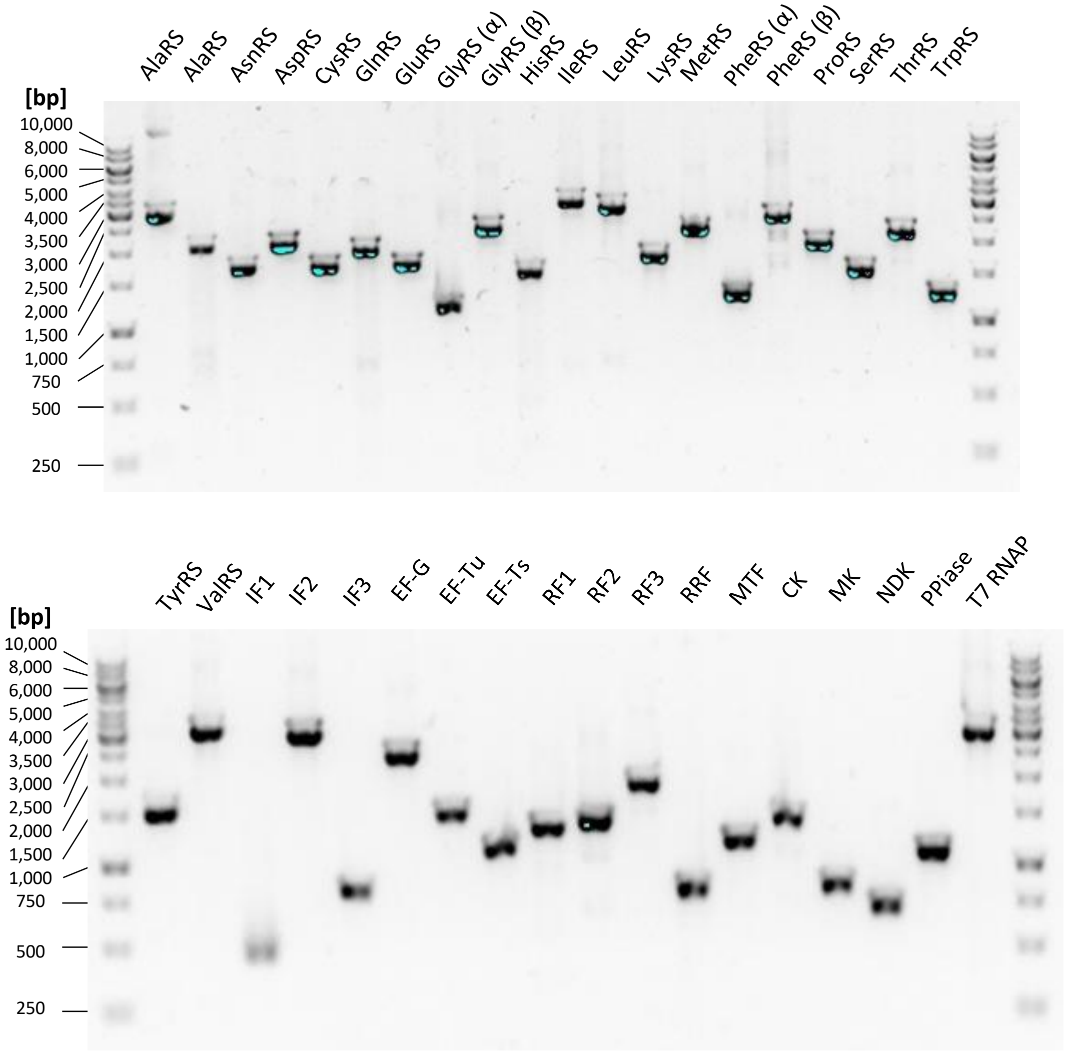


Figure S2: **Amplified and purified linear DNA templates encoding each non-ribosomal PURE protein run on an agarose gel.** For GlyRS and PheRS, which each consist of two subunits, two linear DNA templates were amplified, with each template encoding one subunit.

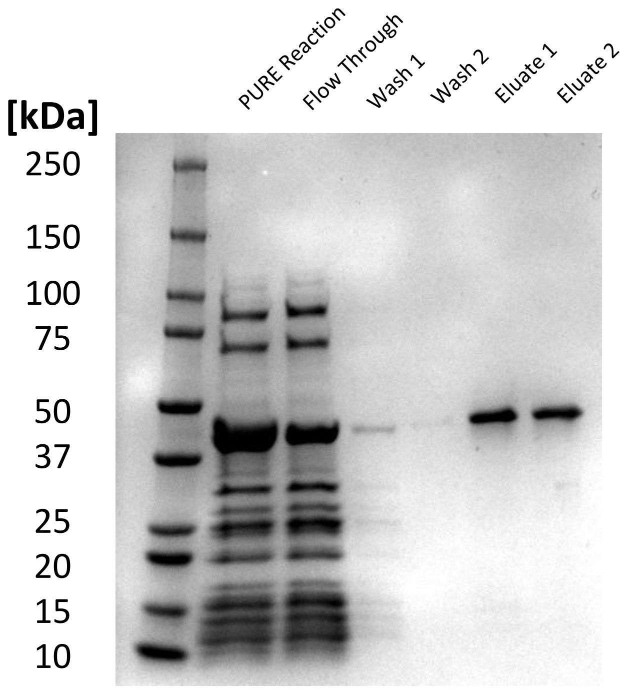


Figure S3: **Coomassie-stained SDS–PAGE of EF-Tu purification.** EF-Tu was expressed in a PURE-frex reaction and purified using magnetic beads. Samples from each purification step were run on SDS–PAGE followed by Coomassie staining.

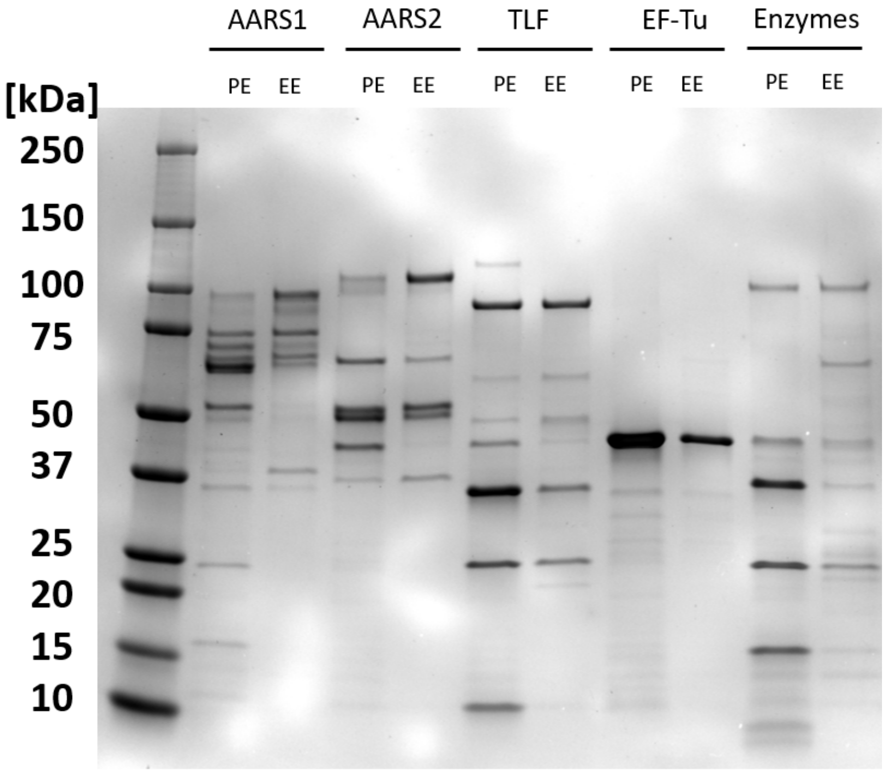


Figure S4: **Coomassie-stained SDS-PAGE of final subsets.** Final subsets, which are labeled as PE, were loaded on SDS-PAGE next to their *E. coli* expressed subsets with adjusted concentration, which is labeled as EE, followed by Coomassie staining. In all lanes, 2 µL was loaded, except for EF-Tu, where 1 µL was loaded. EE: *E. coli* expressed, PE: PURE expressed.

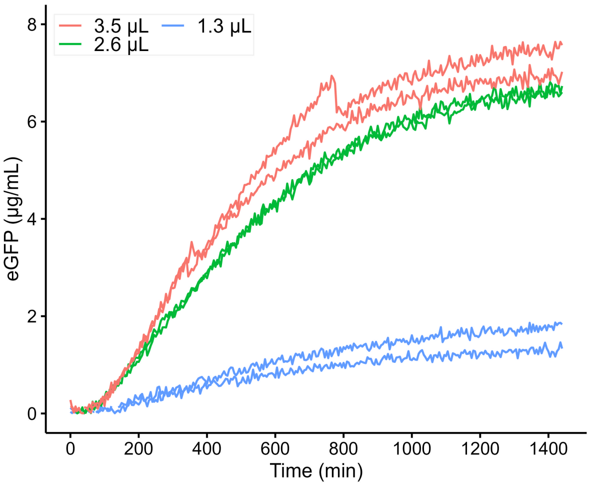


Figure S5: **Titration of diluted full PURE in cell-free reactions.** Different volumes of 20-fold–diluted homemade PURE (containing a 20-fold lower concentration of non-ribosomal proteins) were added to the other reaction components, including the energy solution, ribosomes, and eGFP template, to determine the optimal amount of diluted non-ribosomal PURE protein to supplement the reaction. Fluorescence was measured over time.

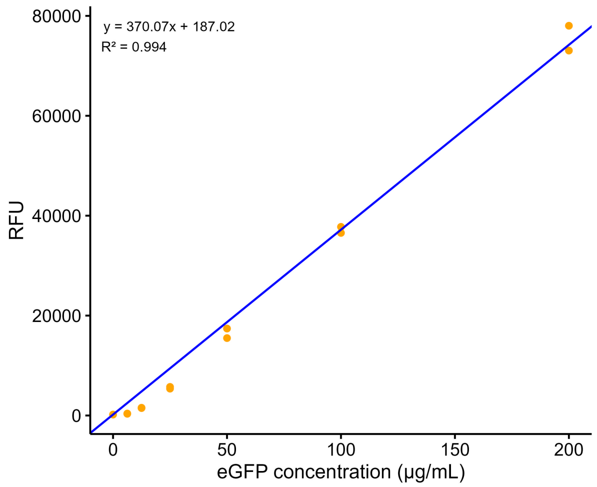


Figure S6: **Calibration curve of eGFP.** Different concentrations of recombinant eGFP were prepared in a PUREfrex reaction master mix lacking a DNA template, and their fluorescence was measured with the same settings as the main samples (excitation: 488 nm; emission: 507 nm; gain: 70%). The average value for the second hour was used as the fluorescence signal. The regression curve was constrained to pass through the mean RFU at 0 eGFP concentration. Each concentration was prepared in duplicate. RFU: relative fluorescence units.

| Indiviudal subset functional tests | | | | |
|---|---|---|---|---|
| Subset | PE subset stock conc. (µg/mL) | PE subset conc. in reaction (µg/mL) | HM PURE conc. in reacion (µg/mL) | PE/HM ratio |
| AARS1 | 412.5 | 41.25 | 194.5 | 4.7 |
| AARS2 | 457 | 45.7 | 158.2 | 3.5 |
| TLF | 394 | 39.4 | 212.8 | 5.4 |
| EF-Tu | 455 | 45.5 | 211.2 | 4.6 |
| Enzymes | 357.5 | 35.75 | 58 | 1.6 |

| Reconstituted PURE functional test | | | | | | |
|---|---|---|---|---|---|---|
| Subset | PE subset stock conc. (µg/mL) | Mixing ratio | Volume in reaction (%) | PE subset conc. in reaction (µg/mL) | HM PURE conc. in reaction (µg/mL) | PE/HM ratio |
| AARS1 | 412.5 | 2.5 | 2.2 | 8.9 | 194.5 | 21.8 |
| AARS2 | 457.0 | 2.5 | 2.2 | 9.9 | 158.2 | 16.0 |
| TLF | 394.0 | 10.0 | 8.7 | 34.1 | 212.8 | 6.2 |
| EF-Tu | 455.0 | 10.0 | 8.7 | 39.4 | 211.2 | 5.4 |
| Enzymes | 357.5 | 5.0 | 4.3 | 15.5 | 58.0 | 3.7 |
| | | | **Sum** | 107.9 | 834.7 | 7.7 |

| Single-reacion regenerated PURE functional test | | | | |
|---|---|---|---|---|
| SRR PURE conc. (µg/mL) | Volume in reaction (%) | SRR PURE conc. in reaction (µg/mL) | HM PURE conc. in reacion (µg/mL) | PE/HM ratio |
| 702.0 | 26.0 | 182.5 | 682.5 | 3.7 |

Table S1: **Concentrations of PE subsets, reconstituted PURE, and SRR PURE compared to their corresponding concentrations in the original PURE formulation.** The top table summarizes the concentrations of PE subsets used for the functional assays in Figure 1d–e. The middle table summarizes the concentrations of subsets and their total concentration in reconstituted PURE (combined PE subsets) used for the functional assays in Figure 2c–d. The bottom table summarizes the concentrations of SRR PURE used for the functional assays in Figure 2h–i. All concentrations were determined by the Bradford assay. PE: PURE expressed, SRR: single-reaction regenerated, HM: homemade.

| | |
|---|---|
| eGFP linear template<br>(Red: T7 promoter,<br>Blue: RBS,<br>Green: eGFP coding sequence,<br>Purple: T7 terminator) | gatcttaaggctagagtactaatacgactcactatagggagaccacaacggtttccctctagaaataattttgtttaacttaag aaggaggaaaaaaaaATGTCTAAAGGTGAAGAATTATTCACTGGTGTTGTCCCAATTTTGGTTGAATTAGATGGTGATGTTAATGGTCACAAATTTTCTGTCTCCGGTGAAGGTGAAGGTGATGCTACTTACGGTAAATTGACCTTAAAATTTATTTGTACTACTGGTAAATTGCCAGTTCCATGGCCAACCTTAGTCACTACTTTAACTTATGGTGTTCAATGTTTTTCTAGATACCCAGATCATATGAAACAACATGACTTTTTCAAGTCTGCCATGCCAGAAGGTTATGTTCAAGAAAGAACTATTTTTTTCAAAGATGACGGTAACTACAAGACCAGAGCTGAAGTCAAGTTTGAAGGTGATACCTTAGTTAATAGAATCGAATTAAAAGGTATTGATTTTAAAGAAGATGGTAACATTTTAGGTCACAAATTGGAATACAACTATAACTCTCACAATGTTTACATCATGGCTGACAAACAAAAGAATGGTATCAAAGTTAACTTCAAAATTAGACACAACATTGAAGATGGTTCTGTTCAATTAGCTGACCATTATCAACAAAATACTCCAATTGGTGATGGTCCAGTCTTGTTACCAGACAACCATTACTTATCCACTCAATCTGCCTTATCCAAAGATCCAAACGAAAAGAGAGACCACATGGTCTTGTTAGAATTTGTTACTGCTGCTGGTATTACCCATGGTATGGATGAATTGTACAAATAAcggctgctaacaaagcccgaaaggaagctgagttggctgctgccaccgctgagcaataactagcataaccccttggggcctctaaacgggtcttgaggggttttttgctgaaaggaggaactatatcc |
| Twist forward primer (used for eGFP template amplification) | CAATCCGCCCTCACTACAACCG |
| Twist reverse primer (used for eGFP template amplification) | TCCCTCATCGACGCCAGAGTAG |
| 5’ sequence of PURE templates<br>(Red: T7 promoter,<br>Blue: RBS) | GATCTTAAGGCTAGAGTACTAATACGACTCACTATAGGGGAATTGTGAGCGGATAACAATTCCCCTCTAGAAATAATTTTGTTTAACTTTAAGAAGGAGATATACAT |
| 3’ sequence of PURE templates<br>(Purple: T7 terminator) | GATCCGGCTGCTAACAAAGCCCGAAAGGAAGCTGAGTTGGCTGCTGCCACCGCTGAGCAATAACTAGCATAACCCCTTGGGGCCTCTAAACGGGTCTTGAGGGGTTTTTT |
| Forward primer for PURE templates | GATCTTAAGGCTAGAGTACTAATACGACTCACTATAGGGGAATTG |
| Reverse primer for PURE templates | AAAAAACCCCTCAAGACCCGTTTAGAG |

Table S2: **DNA sequences.** eGFP linear template, 5’ and 3’ sequences included upstream and downstream of the coding sequence for all PURE proteins, as well as the primers used for amplification.

| Components | Buffer A | Buffer B | Buffer HT | Stock buffer |
|---|---|---|---|---|
| HEPES | 50 mM | 50 mM | 50 mM | 50 mM |
| Ammonium chloride | 1000 mM | - | - | - |
| Magnesium chloride | 10 mM | 10 mM | 10 mM | 10 mM |
| Potassium chloride | - | 100 mM | 100 mM | 100 mM |
| Imidazole (pH=7) | - | 500 mM | - | - |
| Glycerol | - | - | - | 30% (v/v) |
| TCEP | 1 mM | 1 mM | 1 mM | 1 mM |

Table S3: **Protein purification buffers.**
